# Network modelling unravels mechanisms of crosstalk between ethylene and salicylate signalling in potato

**DOI:** 10.1101/214940

**Authors:** Živa Ramšak, Anna Coll, Tjaša Stare, Oren Tzfadia, Špela Baebler, Špela Baebler, Yves Van de Peer, Kristina Gruden

**Affiliations:** National Institute of Biology, Department of Biotechnology and Systems Biology, 1000 Ljubljana, Slovenia; Department of Plant Systems Biology, VIB, 9052 Ghent, Belgium; Department of Plant Biotechnology and Bioinformatics, Ghent University, 9052 Ghent, Belgium; Genomics Research Institute, University of Pretoria, Private bag X20, Pretoria 0028, South Africa

## Abstract

To provide means for novel crop breeding strategies, it is crucial to understand the mechanisms underlying the interaction between plants and their pathogens. Network modelling represents a powerful tool that can unravel properties of complex biological systems. Here, we build on a reliable Arabidopsis (*Arabidopsis thaliana* L.) immune signalling model, extending it with the information from diverse publically available resources. The resulting prior knowledge network (20,012 nodes, 70,091 connections) was then translated to potato (*Solanum tuberosum* L.) and superimposed with an ensemble network inferred from potato time-resolved transcriptomics data. We used different network modelling approaches to generate specific hypotheses of potato immune signalling mechanisms. An interesting finding was the identification of a string of molecular events, illuminating the ethylene pathway modulation of the salicylic acid pathway through NPR1 gene expression. Functional validations confirmed this modulation, thus confirming the potential of our integrative network modelling approach for unravelling molecular mechanisms in complex systems.

**One-sentence summary:** Analysis of integrated prior knowledge and ensemble networks highlights a novel connection between ethylene and salicylic acid signalling modules in potato.

## INTRODUCTION

Plants have evolved a multi-layered immune system to help them cope with potential invasion of pathogens (Jones and Dangl, 2006). Recognition of invading organisms triggers a rapid induction of signalling cascades, leading to diverse defence responses (Pieterse et al., 2012). Effectiveness of these downstream events is crucially dependent on salicylic acid (SA), jasmonic acid (JA) and ethylene (ET), but other hormones were also shown to play important roles in plant immunity (Verma et al., 2016). Hormonal signals differ considerably in timing, quantity and composition, depending on the type of attacker (Bluthgen, 2015). Crosstalk between hormonal pathways can have antagonistic or synergistic effects and is largely multi-dimensional (Tsuda and Somssich, 2015). This interconnected plant hormonal network provides an important regulatory mechanism, granting plants quick adaptation abilities via intruder-specific alterations (Pieterse et al., 2012). At the molecular level, crosstalk between signalling pathways with several regulatory feedback loops adds robustness to the plant immune signalling network (Windram and Denby, 2015). Network modelling, a subfield of systems biology, continues to be paramount in investigations of complex systems (Barabasi, 2009). This approach facilitates thorough analyses of critical system properties and discovery of novel key players or interactions, thus being suitable for providing new insights into plant defence specificities (McCormack et al., 2016).

While heterogeneous technologies of high-content omics allow us to capture snapshots of the systems, the challenge now lies in integration of knowledge into a coherent systems view (Hillmer and Katagiri, 2016). Network inference from omics datasets allows us to deduce the underlying structure of activated processes. However, due to high noisiness, high dimensionality and low sample sizes of data, this is a non-trivial task (Veiga et al., 2010). Thus, additional improvements are needed, for example, incorporation of prior knowledge can greatly improve reconstructed network accuracy, simultaneously reducing noise and sparsity effects of the source data, without inflating the computational cost (Ghanbari et al., 2015).

Despite extensive potato breeding programs, its average yields still do not reach their physiological potential (Singh, 2008). This is the result of its sensitivity to a wide range of environmental factors. The aim of the current study was to improve understanding of potato immune signalling using network modelling and thus on the long-term to provide means for novel crop breeding strategies directed towards high and sustainable yields. We built on a manually curated plant immune signalling model (Miljkovic et al., 2012), complementing it with knowledge from various public resources, the majority of the available for the model plant *Arabidopsis thaliana*. We also inferred networks using time-resolved transcriptomics data of both compatible and incompatible potato-virus interactions (Stare et al., 2015; Baebler et al., 2014) and superimposed them with our knowledge network. We tested the resulting network for its potential for generating novel hypotheses and show that network analysis revealed a previously unknown connection between ET and SA signalling, namely that activation of the ET signalling module regulates expression of NPR1, an important regulator of SA signalling. This newly identified crosstalk was experimentally validated in potato.

## RESULTS

### Construction of the comprehensive knowledge network

Firstly, a previous plant immune signalling model (PIS-v1(Miljkovic et al., 2012)) was expanded with manually curated knowledge from recently published literature. Addition of 64 Arabidopsis genes to the existing model resulted in an expanded PIS-v2 model with 212 biological components (177 genes, 31 metabolites and 4 small RNAs), categorized into 108 component families as defined by Miljkovic et al. (Miljkovic et al., 2012). On the component family abstraction, we added 32 new reactions, to a total 111 reactions (Supplemental Data Set 1).

We combined the graph of binary PIS-v2 interactions with three layers of publically available information: protein-protein interactions (PPI), transcriptional regulation (TR) and regulation through microRNA (miRNA). This resulted in an *Arabidopsis thaliana* comprehensive knowledge network (AtCKN), with 20,012 nodes (19,812 genes, 186 miRNA families, 3 metabolites and 11 viral proteins) and 70,091 connections (Figure 1; Supplemental Table 1). Each data layer covers unique gene or miRNA subsets in the full network, with only six nodes present in all four layers, which supports our layer selection as being well suited for inclusion (Figure 2).

**Figure 1:**
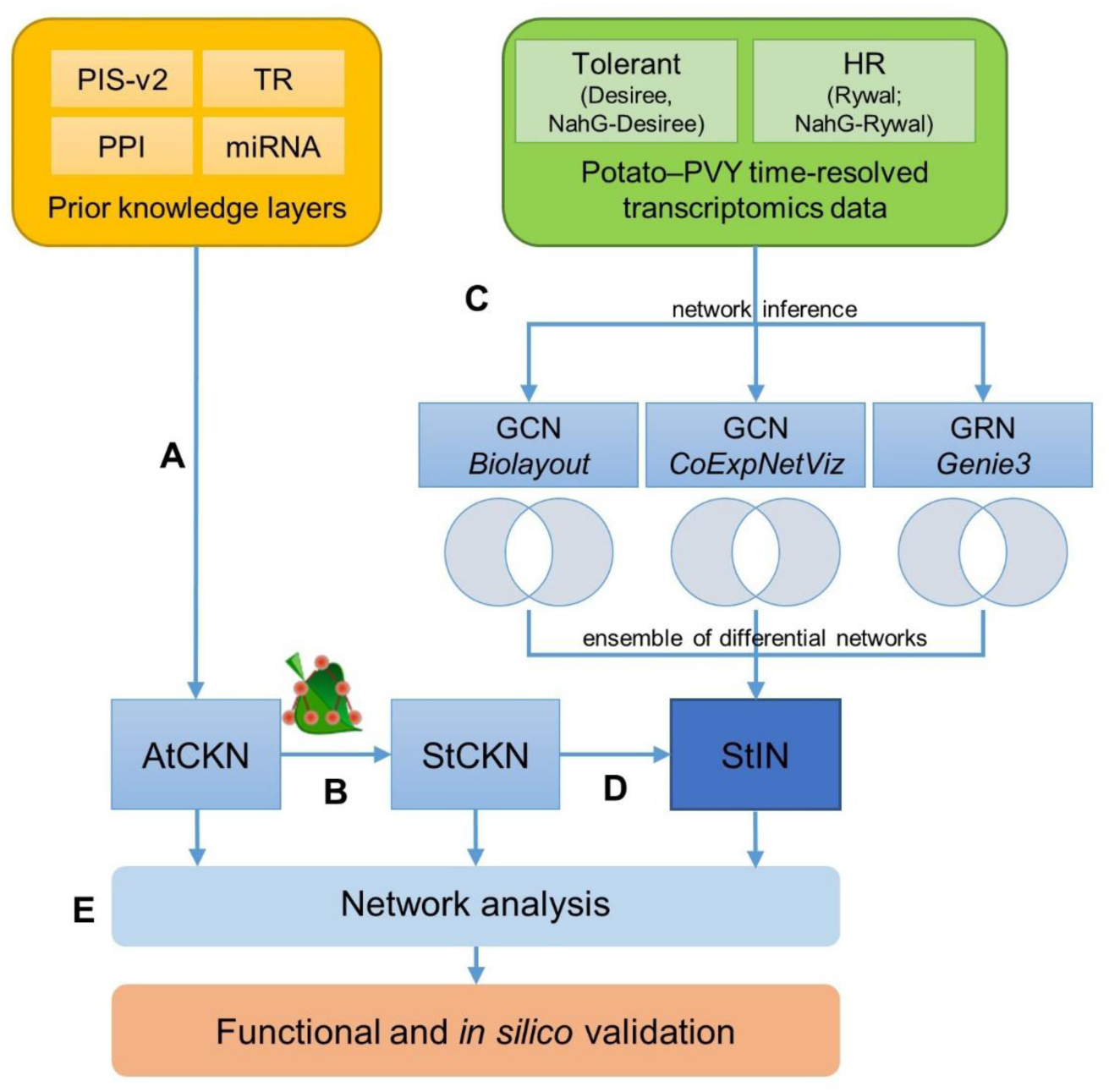
Schematic overview of networks construction and analyses. **a**, Networks of four prior knowledge layers were merged into the *Arabidopsis thaliana* comprehensive knowledge network (AtCKN). **b**, Orthologous relationships were used to translate from Arabidopsis to potato, forming the *Solanum tuberosum* comprehensive knowledge network (StCKN). **c**, Starting with two time-resolved transcriptome datasets, gene co-expression networks (GCN; relationships between co-expressed genes) and gene regulatory networks (GRN; transcription factor to regulated gene relationships) were inferred using three methods. For each inference method, two subnetworks were generated for mock-inoculated and viral-infected samples. Removing all connections present in all co-expression or gene regulatory networks resulted in differential networks, respectively. **d**, StCKN and differential networks were merged into the potato integrated network (StIN). **e**, Created networks were analysed using network analysis approaches. PIS-v2 –plant immune signalling model; PPI – protein-protein interactions; TR – transcriptional regulation; miRNA – regulation via miRNA.

**Figure 2:**
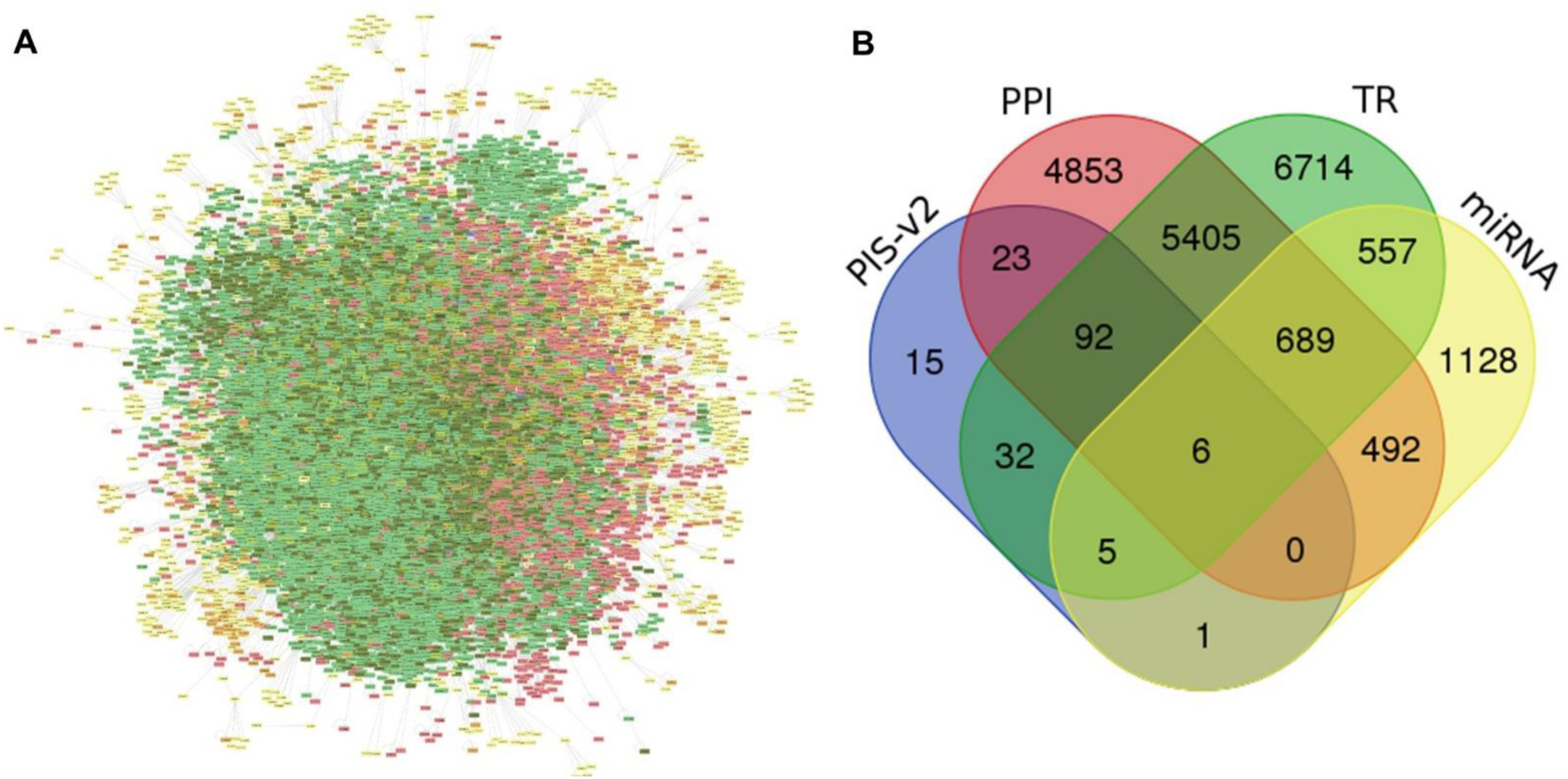
Contribution of the four layers to node coverage in *Arabidopsis thaliana* comprehensive knowledge network (AtCKN). **a**, Prefuse force directed layout of the largest connected component in AtCKN. **b**, Display of overlap between the four contributing layers to AtCKN. PIS-v2 – plant immune signalling model; PPI – protein-protein interactions; TR – transcriptional regulation; miRNA – regulation via miRNA. Different colors represent the same entities in both panels.

### Using prior knowledge to improve the plant immune signalling model

To assess the potential of using prior knowledge for the improvement of the mechanistic model of plant immune signalling, we extracted a subnetwork of AtCKN with all components of PIS-v2. This subnetwork consisted of 391 connections between 212 nodes in the fully expanded version or 254 connections between 108 nodes at the level of component families. By comparing the PIS-v2 layer against the remaining layers of AtCKN (PPI, TR, miRNA), we found 45 connections present in both subnetworks, 67 were present only in the PIS-v2, and 142 novel reactions from the remaining AtCKN. These represent model upgrades, demonstrating the value of dispersed knowledge sources also for the construction of detailed mechanistic models. Inspecting these new connections showed that manual curation is more successful in knowledge extraction within a signalling module than between signalling modules (Figure 3; Supplemental Data Set 2).

**Figure 3:**
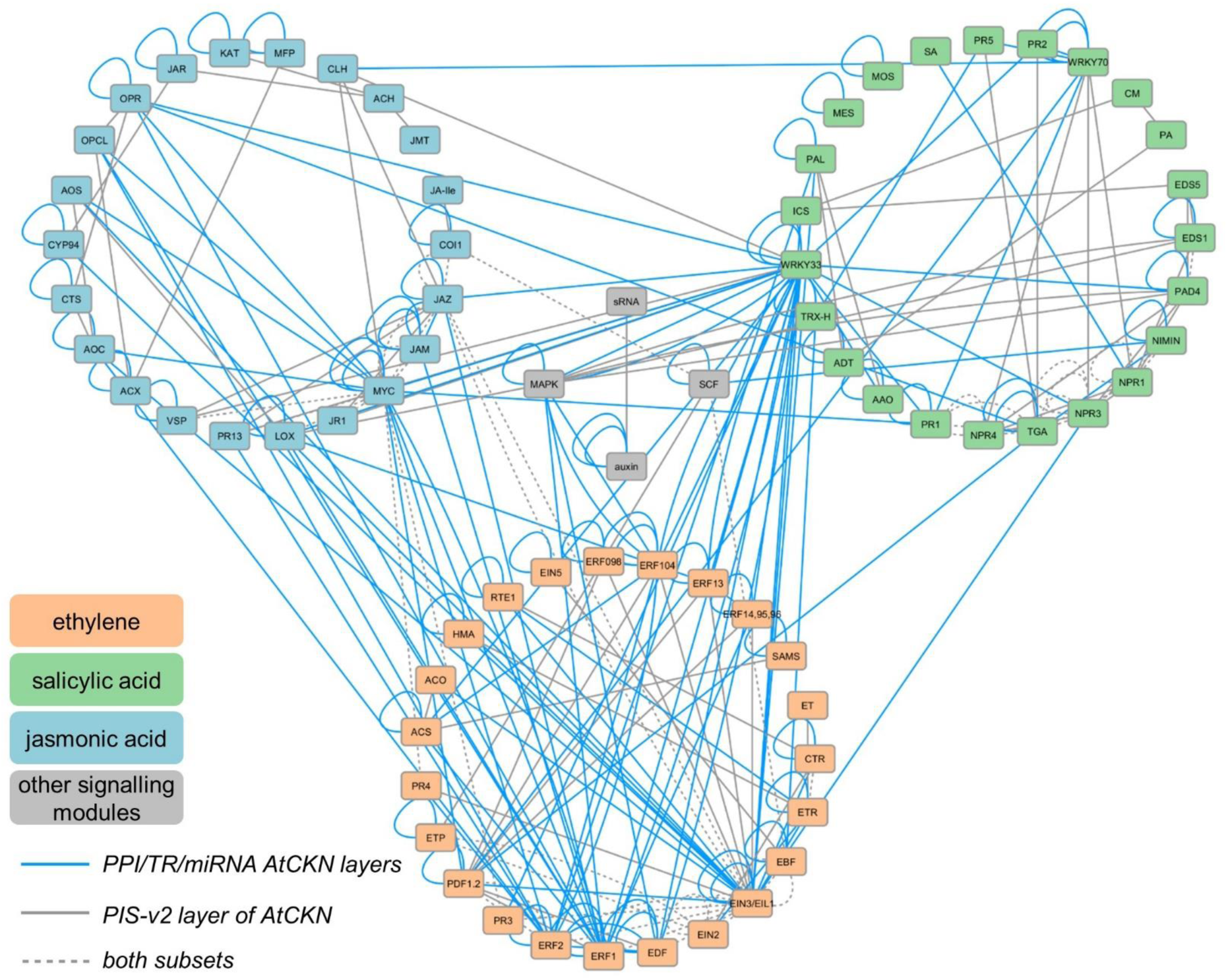
Contribution of PIS-v2 layer in contrast to the remaining AtCKN layers (PPI, TR, miRNA). Connections between gene families of three plant hormone immune signalling pathways (ET – orange, JA – blue, SA – green) and other signalling modules (grey). Blue full lines – novel connections present in PPI, TR and miRNA layers of AtCKN; grey full lines – connections present only in the PIS-v2 layer; dotted grey lines – connections existing in both compared subsets.

### Translation of knowledge to potato and integration with experimental data

Based on predictions of orthologous relationships, we translated AtCKN from Arabidopsis to potato. As expected, finding orthologous genes between plant species is often delineated as a many-to-many relationship, as well as cases where no homologous gene is found. This resulted in an intermediary abstracted network with 9679 nodes (9497 orthologue groups, 168 miRNA families, 3 metabolites, 11 viral proteins) and 43,393 connections. Next, we inferred all combinations between potato genes of the same orthologue group for each abstracted connection, resulting in StCKN with 18,036 nodes (17,855 genes, 168 miRNA families, 3 metabolites, 11 viral proteins) and 296,834 connections (Figure 1).

To identify transcriptional modules in potato that contribute to potato immune signalling in PVY infection, we selected datasets profiling temporal response dynamics in potato genotypes displaying either a tolerant or hypersensitive response (108 samples). Out of 17,855 potato StCKN genes, 10,920 (61%) had a microarray probe assigned. We used the expression values of these genes to infer a targeted and non-targeted co-expression network and a gene regulatory network: the former in order to propose genes controlled by the same transcriptional regulatory program, and the latter to propose potential regulators (Figure 1).

Two types of subnetworks generated (mock-inoculated and viral-infected) allowed us to examine differences between gene connections. Subnetworks of mock-inoculated plants reflecting developmental cues were all of similar size (25,916, 25,910 and 30,570 connections for targeted, non-targeted co-expression and gene regulatory network, respectively). Sizes of subnetworks reflecting plant responses to viral infection were between 56-64% of mock-inoculated subnetworks, but again similar when compared to each other (16,716, 15,993 and 17,204 connections in the same order as above). The difference in connection count per each method could be explained by a greater variability of the viral subset (tolerant, hypersensitive), resulting in a smaller number of inferred connections. Conversely, as developmental profiles of all genotypes share greater similarities, the number of inferred connections was larger in the mock-inoculated subnetwork. Comparison of predicted connections among all three approaches revealed they are largely independent, as the three methods shared only 111 out of 116,391 total unique connections (Supplemental Figure 1).

To elucidate perturbations in network topologies in plant immunity, we extracted differential networks by removing any connections shared between the mock-inoculated and viral-infected treatments. The differential networks were merged as new layers with StCKN into a potato integrated network (StIN; Figure 1) with 402,277 connections between 19,801 nodes (19,619 genes, 168 miRNA families, 3 metabolites, 11 viral proteins).

To validate our approach of StIN construction, we compared interactions covered by selected layers of biological information against a gold-standard, a set of highly reliable reactions from the manually curated plant immune signalling model (Table 1). The PPI layer covered 50% of all reactions identified by manual literature curation in our PIS-v2 model. The TR layer had even greater concordance with the gold-standard reactions (80%). On the other hand, connections resulting from gene regulatory network inference covered only 20% of interactions in the gold-standard. We must however note that some PIS-v2 model connections might be triggered in non-viral infections instead.

**Table 1:**
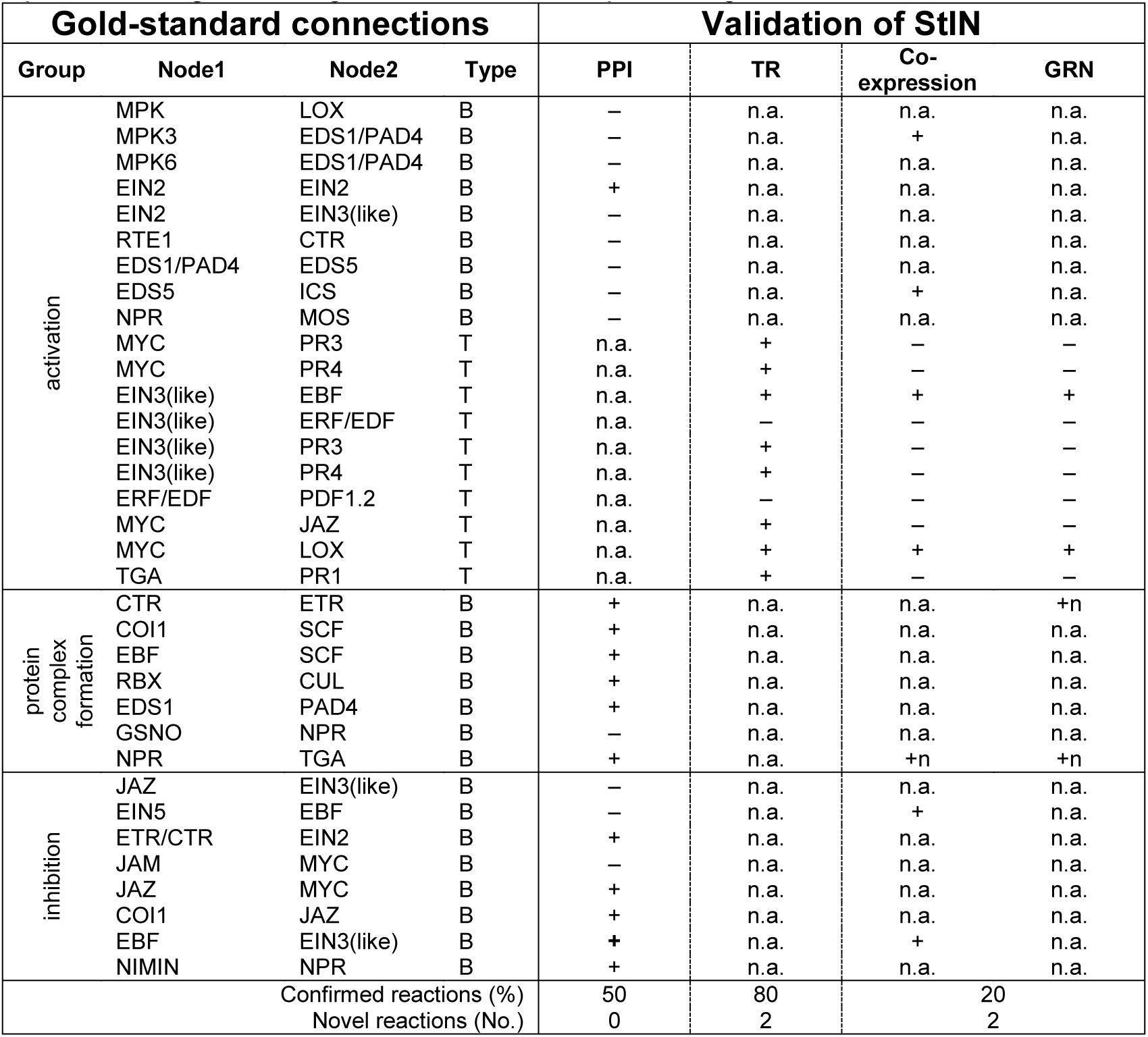
StIN validation by comparing connection existence in selected data layers against the gold-standard. Connections grouped on their reaction effect types (activation, binding, inhibition) and their interaction type (B – protein binding, T – transcriptional regulation). Compared layers include partial StCKN (PPI, TR) and two differential networks (merged targeted and non-targeted co-expression and gene regulatory). Coverage of a gold-standard reaction (on the component family level) is indicated by + (reaction present in the layer), +n (reaction present in the layer, but not in the gold-standard), – (reaction not present in the layer) or n.a. (not a relevant comparison – e.g. transcriptional regulation can’t validate a protein binding connection). Co-expression results were not included in validation as they represent co-regulation of genes and not transcriptional regulation.

### Integrated network-driven hypotheses: Ethylene is modulating NPR1 gene expression

First, we analysed the topologies of the generated networks, namely AtCKN, StCKN and StIN. AtCKN showed some bias towards high degree nodes, a direct result of two included datasets from ChIP-Seq experimental data (Supplemental Table 1). On the other hand, expansion to all potato genes performed for StCKN and StIN distorted the network topological indices (many-to-many phylogenetic relationships). Further network analyses aimed at targeted identification of novel crosstalk connections, between receptors and transmitters of seven plant hormonal pathways (Supplemental Table 2). Due to topology distortion in translated potato networks, we performed the initial search in AtCKN and afterwards analysing them in StCKN.

One of the most interesting findings was the shortest path from the ethylene pathway transmitter EIN3 to SA receptor NPR1. In AtCKN we identified several shortest paths of 3-step length, involving 32 genes and 61 connections (Figure 4). From those, one third were binding (PPI) connections and the remainder transcriptional regulations. In StCKN, we searched for walks (length 3) from EIN3 to NPR1, and then superimposed co-expression and GRN connections from StIN (Figure 4). The potato EIN3 to NPR1 walk subnetwork included 32 genes, 57 StCKN and 48 experimentally inferred connections. Searching for walks of a specific length was required to ease comparisons, as the shortest path between EIN3 and NPR1 after translation was of 2-step length (Figure 4).

**Figure 4:**
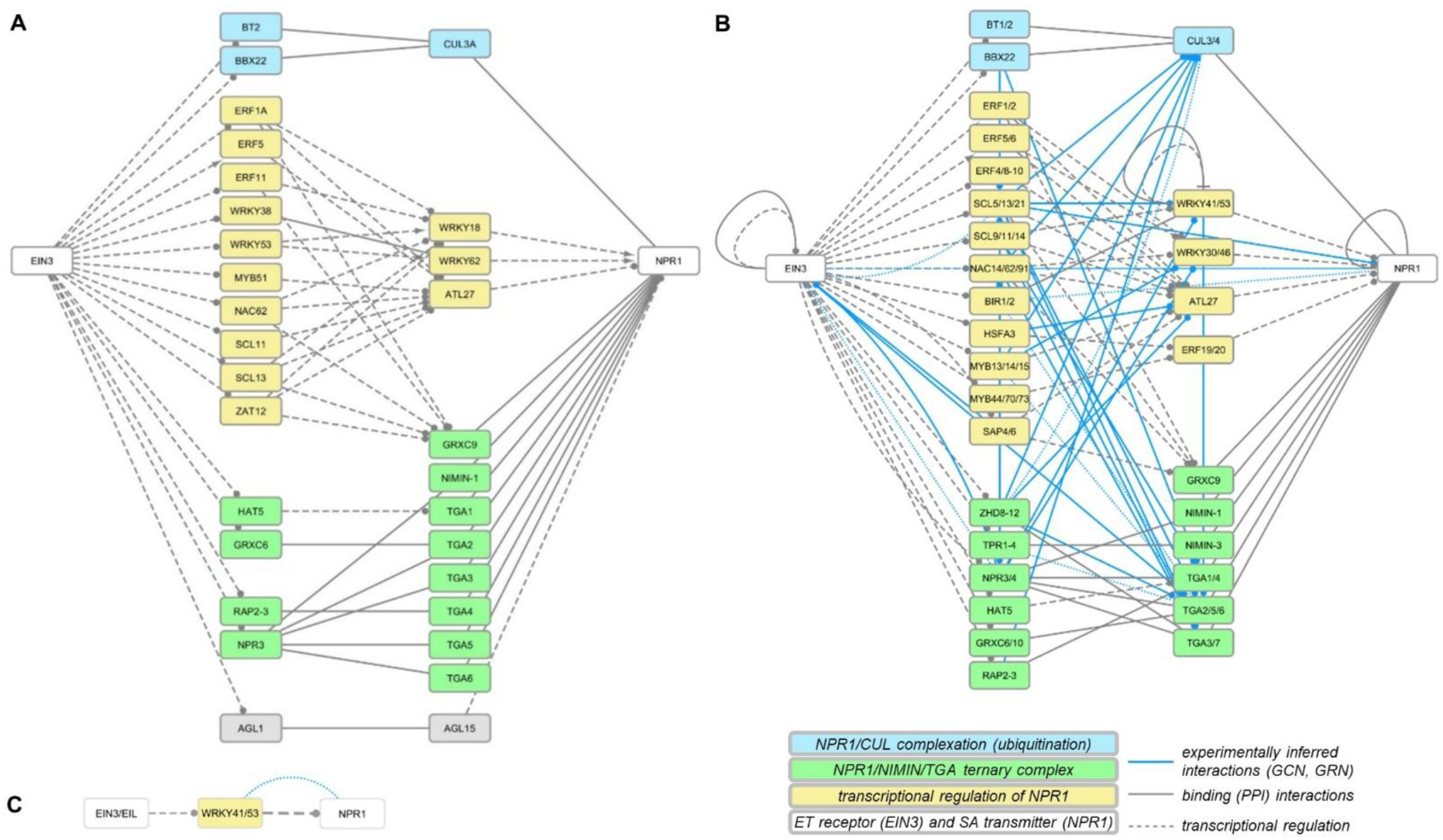
Results of shortest paths or walks from EIN3 (ET) towards NPR1 (SA). **a**, Shortest path (3-steps) in AtCKN; **b**, walk of length 3 in StCKN; **c**, shortest path of (2-steps) in StCKN. Line type and colour indicate the interaction type: binding (grey solid), regulation by transcription factors (grey dash), co-expression (blue dots) and gene regulatory (blue solid). Target arrow indicates the action of the connection: activatory (arrow), inhibitory (T), unknown (circle) or undirected in the case of binding or co-expression (no arrow). Gene group identifiers corresponding to these images are given in Supplemental Data Set 3.

The majority of the binding type connections in potato EIN3-NPR1 walk can be attributed to the formation of a ternary complex between NPR1, NIMIN1 and TGA factors (Figure 4, green), which in turn modulates PR1 gene expression (Weigel et al., 2005). The other set of binding type connections relates to complexation of NPR1 with cullin (Figure 4, blue), which is important for plant immunity regulation (Spoel et al., 2009). The remaining shortest paths indicate potential transcriptional regulation of NPR1 through ET signalling (Figure 4, yellow). As expected, superimposed connections from transcriptomics potato data were also denser in this area (Figure 4, blue connections). The first step of all shortest paths describing transcriptional regulation includes several ethylene responsive factors (ERF) and specific members of the C2C2, GRAS, NAC, MYB and HSF families. These transcription factors then target two WRKYs, in particular WRKY18 and WRKY62, where the former was shown to have an important role in plant responses to bacterial and fungal pathogens (Chen et al., 2010; Xu et al., 2006), besides another ERF and an ATL.

Superimposing inferred connections based on transcriptional profiles of potato response to viral infection, although incomplete as they are, confirmed potential regulation of NPR1 gene activity through several WRKY (WRKY30/46, WRKY41/53), zinc-finger (SAP4/6) and MYB (MYB13/14/15, MYB44/70/73) transcription factors (Figure 4). Interestingly, when searching in a reduced StIN with only transcriptional layer connections, we predicted regulation of NPR1 through ERF (ERF2a), WRKY (WRKY34, WRKY41/53), MYB (MYB18/19, MYB52/54) and bHLH (bHLH84/84) transcription factors (Supplemental Figure 2).

For further evaluation of these findings, we scanned Arabidopsis and potato promoters of NPR1 gene for known *cis*-regulatory elements. Apart from containing known conserved motifs for light and development responses, both promoters contain motifs specific for several hormones (abscisic acid, gibberellic acid, JA and SA; Supplemental Data Set 4). In terms of general stress responses, both promoter regions contain binding sites for responses to heat, drought and defence. In addition, we detected a wounding motif, MYB and WRKY binding motifs in the potato promoter only.

### Experimental validation of transcriptional regulation of NPR1 by ET

To validate our network-generated hypothesis, we tested the transcriptional regulation of NPR1 following the induction of ET pathway in potato as more data was supporting our hypothesis for this plant. We additionally checked for the potential of SA signalling module to participate in this process. Thus, we induced the SA signalling module by INA (a functional analogue of SA that is not accessible to degradation by NahG) while either leaving the ET module active or blocking its activity (treatment with 1-MCP). Alternatively, we have tested the regulatory potential of ET module, while SA signalling was blocked by using transgenic plants expressing NahG, which degrade any internally produced SA (Figure 5). Induction of ACO4 gene expression was used as a marker of efficient activation of ET signalling module and PR1b as marker of SA signalling module activation.

**Figure 5:**
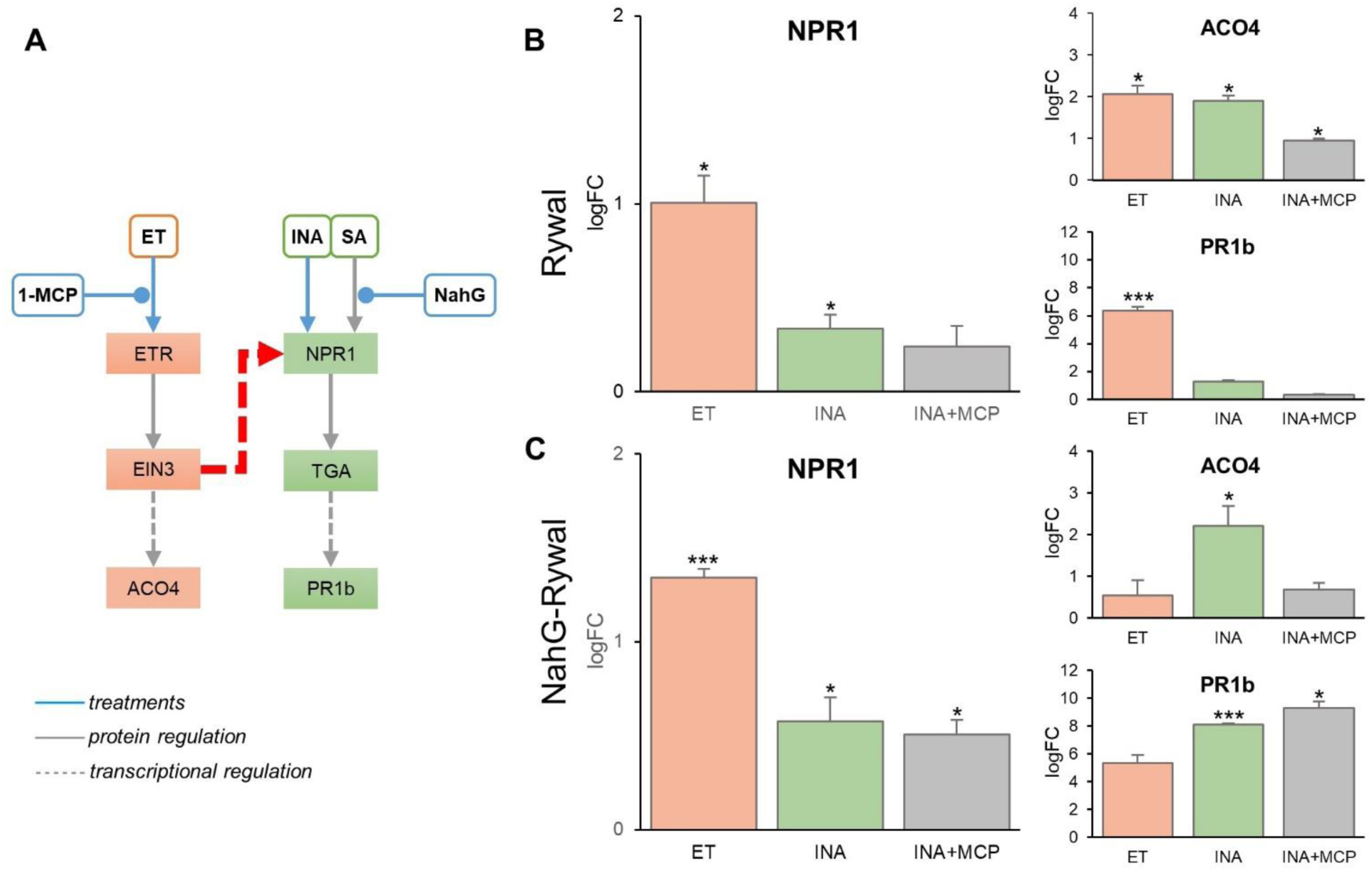
Validation of direct transcriptional regulation of NPR1 by the ethylene (ET) signalling module in potato leaves. **a**, Schema of the underlying biological pathways of ET (orange) and salicylic acid (SA; green), with interactions on the protein or transcriptional level (solid and dashed lines, respectively). Interactions can activate (arrow) or inhibit (circle) downstream signalling events. Blue coloured lines denote treatments of this experiment (ET, INA, 1-MCP) or NahG plants (deficient in SA signalling). The red dashed arrow denotes our tested hypothesis. **b**, Potato plants cv. Rywal and **c**, its transgenic line NahG-Rywal were treated with either ET (orange), INA (SA analogue; green) or a combination of INA with 1-MCP (ET inhibitor; INA+MCP; grey). Log2 fold change in gene expression of treated versus control plants is shown (*-p<0.05, **<0.01, ***<005; n=3) for ACO4 (ET signalling marker), PR1b (SA signalling marker) and NPR1 (SA signal transmitter). Error bars denote standard error of biological replicates. Note different y-axis scales for different genes. Results of the second independent experiment are in Supplemental Figure 3.

Significant upregulation of NPR1 gene expression after ET treatment substantiated our network-generated hypothesis in potato plants (Figure 5). Strong induction of the PR1b gene after ET treatment additionally confirmed regulation of the SA signalling module by ET. Induction of PR1b by ET was even stronger than its expected and well-established induction by SA signalling (Figure 5, INA treatment). Tight interaction between both modules was also confirmed by ACO4 induction by both ET and INA treatment. When blocking SA signalling (using NahG transgenic plants, Figure 5), the induction of NPR1 gene expression was similar as in non-transgenic plants, confirming that the string of events leading to activation is not dependent on SA. All other module crosstalk observed in non-transgenic plants was also confirmed in the SA-depleted plants.

## DISCUSSION

Plants have evolved a complex immune system to defend themselves against diverse pathogens and herbivores. This plant immune signalling network with its tightly interconnected signalling modules ensures a timely, precise and effective response to attackers (Coolen et al., 2016). Many regulatory mechanisms are buffered by the network, making them undetectable by traditional genetic approaches of single-gene null-mutant analyses (Hillmer et al., 2017), making network modelling algorithms invaluable tools to expand our understanding of plant immunity (Windram and Denby, 2015).

An ideal network model would encompass all components of the biological system and all interactions between them. However, due to limited knowledge and data availability, this is currently not possible. To circumvent this problem, researchers studying plant immune signalling have taken various approaches. Bottom-up approaches based on manual literature curation lead to detailed and accurate models (Naseem et al., 2012; Miljkovic et al., 2012), but their extensiveness is limited. However, the majority of published research builds on networks inferred entirely from experimental data (e.g. (Ebrahim et al., 2016; Vermeirssen et al., 2014)), or a combination of network inference with some kind of prior knowledge (e.g. (Jiang et al., 2016; Sabaghian et al., 2015)), which can significantly simplify the computational burden of network inference (Windram and Denby, 2015).

In this study, compared to other efforts of prior knowledge integration (Dai et al., 2016), we have based our knowledge network on the manually built, highly reliable model of plant immune signalling (Miljkovic et al., 2012) and complemented it with knowledge from various databases or datasets published as supplements of manuscripts in majority available for the Arabidopsis. Therefore, our results represent the currently most comprehensive knowledge network of immune signalling and related processes in Arabidopsis (see Data availability section of Methods). Compared to this model species, data on immune signalling is much lesser for crop species, therefore the translation of knowledge represents a basis for crop resistance breeding. We have translated the AtCKN to StCKN and subsequently inferred networks from time-resolved experimental data on potato–virus interaction. To alleviate problems of unknowns in the dynamics of potato response to PVY, we applied different methods to construct both co-expression and gene regulatory networks (Figure 1). Such ensemble solutions have been shown to match or outperform single methods, particularly in revealing the true underlying network structure (Vermeirssen et al., 2014).

To assess the validity of our approach, we compared interactions covered by different layers of biological information against a gold-standard, i.e. a set of reactions from the manually curated plant immune signalling model (Table 1). We show a coverage of 58% of known interactions in the newly built network. Biologically more relevant, our integrated network approach predicted 142 additional connections between components of the manually built PIS model, which shows the potential of our integrated network approach in generating new testable hypotheses about biological systems (Figure 3).

As the comparison of the PIS model with other knowledge sources revealed low coverage of connections between signalling modules in the literature (Amar and Shamir, 2014) (Figure 3), we tested the power of our newly built model to identify crosstalk connections. We have indeed discovered an interesting novel transcriptional regulation connection between ET and SA signalling modules, where EIN3 regulates the transcription of NPR1 gene (Figure 4). Further, we confirmed this regulatory connection both by *in silico* promoter analysis (Supplemental Data Set 4) and a set of experiments in potato (Figure 5; Supplemental Figure 3).

Evidence on the importance of ET in plant immune signalling is emerging from several perspectives (Broekgaarden et al., 2015). For example, ET biosynthesis was found to be crucial for induction of programmed cell death during interaction of *Nicotiana umbratica* with *Alternaria alternata* (Mase et al., 2012) and *Pseudomonas syringae*-triggered Arabidopsis susceptibility to herbivory (Groen et al., 2013). Several experiments have shown that SA can modulate ET signalling (Caarls et al., 2016; Guan et al., 2015; Zander et al., 2014; Van der Does et al., 2013). This regulation is mostly implicated in the context of JA/SA antagonism (Caarls et al., 2015; Derksen et al., 2013; Robert-Seilaniantz et al., 2011). There are only a few examples of studies indicating that ET might be an important regulator of SA signalling. It was shown that EIN3 transcription factors directly target the promoter of ICS2, negatively regulating SA biosynthesis (Chen et al., 2009). On the other hand, Frye et al., Mikkelsen and Leon-Reyes et al. (Frye et al., 2001; Mikkelsen, 2003; Leon-Reyes et al., 2009) have shown ET potentiation of PR1 gene expression in Arabidopsis. We observed the same effects in our potato experiments (Figure 5). Chromatin immunoprecipitation sequencing of EIN3 targets revealed NPR3 promoter as its direct target (Chang et al., 2013), providing further evidence that the two modules should be connected.

Most studies involving NPR1 as a master regulator of SA signalling have focused on its posttranslational modifications or loss of function effects. ET modulation of the NPR1 role in JA/SA antagonism was studied using a series of experiments with the *npr1* mutant, which did not allow to discern the effects of transcriptional, translational and posttranslational regulation (Leon-Reyes et al., 2009). Further studies showed the importance of proteasome-mediated degradation of NPR1 (Ding et al., 2016; Saleh et al., 2015; Fu et al., 2012) and nuclear import (Fu et al., 2012; Lee et al., 2015; Kovacs et al., 2015) for effective SA perception. To our knowledge, no study performed so far focused on NPR1 gene expression.

Detailed inspection of our integrated network shows several potential transcription regulation paths from EIN3 to NPR1 gene (Figure 4), involving a cascade of one or two transcription factors. Some of the transcription factors are better characterised (ERF and WRKY), while some were not yet investigated in detail (ATL27 or bHLH), at least not in relation to immune signalling (MYB). As our *in silico* promoter analyses identified WRKY and MYB transcription factor binding motifs, these are the most likely candidates for signal transduction. Considering different experiments, including ours, we reason, that apart from regulation of NPR1 activity on the protein level, transcriptional regulation of NPR1 gene also contributes to immune signalling in plants.

## CONCLUSIONS

We conclude that integration of prior knowledge and experimental datasets followed by network modelling is useful for hypothesis generation, as suggested previously (Medeiros et al., 2015). We also demonstrate its importance in building mechanistic models, which allow for more detailed dynamic modelling of processes. It will also improve mechanistic modelling of plant responses in the multiway interactions, resembling field conditions. By adding effectors, MAMPs of different microorganisms, DAMPS and known triggering points of different abiotic stresses to the network, we can through relatively simple network modelling, understand the mechanisms underlying emerging properties of such complex systems. Additionally, as the prior knowledge network is problem agnostic, the developed approach can also be applied to any other research question in plant biology.

## METHODS

### Network-based knowledge integration

First, a previously inferred plant immune signalling model (Miljkovic et al., 2012) was upgraded by adding manually curated reactions from the recent literature (forming PIS-v2). Further, we transformed the model reactions to a graph of binary interactions, forming the first knowledge layer. Next, we retrieved additional binary connections from different public resources representing additional knowledge layers (full descriptions in Supplemental Table 1); protein-protein interactions (PPI) from databases AtPIN and STRING-v10, two yeast two-hybrid experiments and three experiments on plant-pathogen interactions; transcriptional regulation (TR) from atRegNet and ATRM, two ChIP-Seq experiments and one predicted dataset; regulation through miRNA (miRNA) from miRTarBase, PMRD and PNRD.

These prior knowledge layers were integrated into the *Arabidopsis thaliana* comprehensive knowledge network (AtCKN; Figure 1). Reliability ranks were assigned to each connection between two nodes in AtCKN (Supplemental Table 1). To translate the network from Arabidopsis to potato, a union of three orthologue clusterings was used (available in the GoMapMan database as OCD_all (Ramšak et al., 2014); www.gomapman.org/exports/). Only connections where both Arabidopsis nodes had a defined orthologue in potato were kept in the *Solanum tuberosum* comprehensive knowledge network (StCKN; Figure 1).

### Network inference from experimental datasets

Two microarray datasets profiling temporal response dynamics in potato genotypes with a tolerant (GEO:GSE58593; (Stare et al., 2015)) or hypersensitive (GEO:GSE46180, (Baebler et al., 2014)) response to viral infection were used. Microarray features (microarray probes) were translated to potato gene models (Ramšak et al., 2014). For potato genes covered by several microarray features, one was selected as representative microarray feature based on the maximum number of differentially expressed time points per feature between mock-inoculated and viral-infected samples (FDR corrected p-values <0.05). In case of several such features, the ones with highest log fold change (logFC) and highest average expression across time points were prioritized.

Three network inference methods were applied to the gene expression data. Non-targeted co-expression networks were inferred with BioLayout (Theocharidis et al., 2009), afterwards running the Markov clustering algorithm (Van Dongen, 2008) to divide the graph into discrete subsets. Pearson correlation coefficients (PCCs) were calculated on gene expression of representative microarray features for 156 PIS-v2 potato genes and their 2548 first neighbours in StCKN (PCC ≥ 0.98, top 1 percentile). Targeted co-expression networks were inferred with CoExpNetViz (Tzfadia et al., 2016), calculating co-expression values using mutual information and PCCs (percentiles set between 1 and 99). As bait, 156 potato genes of PIS-v2 were used against all 17,171 representative microarray features. Inference with Genie3 (Huynh-Thu et al., 2010) was performed on the same subset of microarray features as for non-targeted co-expression network inference (weight ≤ 6×10^−3^, top 1 percentile). Thresholds for BioLayout and GENIE3 were determined empirically, so that the resulting networks followed the scale-free and small-world properties of complex networks. Each method was used to generate a mock-inoculated and a viral-infected subnetwork, from data for 16 biological samples each (Figure 1). By removing any connections shared between networks from both treatments (mock-inoculated and viral-infected), a differential network was created for each inference type. Finally, binary interactions in StCKN and both differential networks were merged to create the *Solanum tuberosum* integrated network (StIN).

### Validation of the network construction approach

To assess and estimate the importance and contributions of various knowledge layers in StIN, a gold-standard (set of reliable connections) was constructed from manually curated PIS-v2. Genes were grouped into so-called component families (see Miljkovic et al. (Miljkovic et al., 2012) for representation levels), and connections compared at this level of abstraction. We kept 37 reactions, where all components had both an Arabidopsis and a potato orthologue. StCKN prior knowledge layers (PPI, TR, miRNA) and differential networks were then compared against the gold-standard.

### Network analyses

NetworkAnalyzer (Doncheva et al., 2012) was applied to calculate graph indices and MCODE (Bader and Hogue, 2003) to search for highly interconnected subgraphs in the constructed networks. Pajek (Batagelj and Mrvar, 1998) was applied to search for shortest paths and walks between all combinations of 14 manually selected genes known to be involved in plant signalling (specifically receptors and transmitters for seven plant hormones (Supplemental Table 2). For all network visualisations, Cytoscape (Shannon et al., 2003) was used.

### *In silico* promoter analyses

Sequences 1500 nt upstream of the NPR1 gene translation start site were extracted for Arabidopsis (AT1G64280; Araport 11) and potato (Sotub07g011600.1.1; SpudDB) and scanned for known *cis*-regulatory elements with TRANSFAC (Matys, 2003) and PlantCARE (Lescot et al., 2002).

### Plant growth and treatments

Potato (*Solanum tuberosum* L.) cv. Rywal and its transgenic line NahG-Rywal deficient in SA accumulation (expressing SA hydroxylase; (Baebler et al., 2014)) were cultivated as previously described (Baebler et al., 2009). Treatments were performed on 4-week-old plants. For SA treatments, plants were sprayed with 300 µM INA (2,6-dichloroisonicotinic acid, Aldrich) dissolved in ethanol; control plants were sprayed with 1% ethanol solution. For ET treatments, plants, sealed in airtight clear plastic containers, were treated with 50 ppm ET (Messer); control plants were sealed in identical containers without ET. To inhibit the ET signalling pathway, plants were first treated with SmartFresh (containing 0.14% 1-Methylcyclopropene, 1-MCP; AgroFresh, Inc.) according to manufacturer’s protocol; followed by INA and 1-MCP treatment after 2 h. Potato leaves were sampled 24 h after treatment and immediately frozen in liquid nitrogen (3 plants per each treatment and genotype). The experiment was performed twice.

### Gene expression analysis

Leaf samples (~100 mg) were homogenized with the FastPrep Instrument (MP Biomedicals). Total RNA extraction, DNAse treatment, RNA quality control and reverse transcription were performed as previously described (Baebler et al., 2009). Gene expression was measured using high-throughput qPCR for nonexpresser of PR genes 1 (NPR1), pathogenesis related protein 1b (PR1b) and aminocyclopropanecarboxylate oxidase 4 (ACO4). Cytochrome oxidase (COX) and elongation factor 1 (EF-1) genes were used as endogenous controls (Supplemental Table 3). TATAA PreAmp GrandMaster^®^ Mix (TATAA Biocenter AB) was used for cDNA pre-amplification (2 dilutions per sample) according to manufacturer’s specifications. Gene expression analysis of the samples was held in Fluidigm BioMark™ HD System Real-Time PCR (Fluidigm) using 48.48 Dynamic Arrays IFC. The sample reaction mix contained pre-amplified sample DNA (10-fold diluted), DNA Sample Loading Reagent (Fluidigm) and FastStart Universal Probe Master (Rox; Roche). The assay reaction mix included the Assay Loading Reagent (Fluidigm) and a mix of 2.5 µM TaqMan probe and 9 µM forward and reverse primers. IFC Controller (Fluidigm) was used to prime and load the IFC according to manufacturer’s protocol and under standard PRC reaction conditions. For relative gene expression quantification using standard curve, quantGenius (Baebler et al., 2017); http://quantgenius.nib.si) was used. To determine differences in gene expression between treated and control sample groups, Student’s t-test was performed.

## Data Availability

Microarray transcriptomics data is available from Gene Expression Omnibus (GSE58593, GSE46180). AtCKN network is available from NDEx (Pratt et al., 2015) at this link: http://www.ndexbio.org/#/network/67507c30-995f-11e7-a10d-0ac135e8bacf?accesskey=.

### Supplemental Data

Supplemental Figure 1: Comparison of predicted connections for three selected network inference algorithms.

Supplemental Figure 2: Shortest path search from EIN3 to NPR1 in Solanum tuberosum integrated network (StIN).

Supplemental Figure 3: Results of the replicated experiment for validation of direct transcriptional regulation of NPR1 by the ethylene (ET) signalling module in potato leaves.

Supplemental Table 1: Contribution of four knowledge layers to the built Arabidopsis thaliana comprehensive knowledge network (AtCKN).

Supplemental Table 2: List of selected plant hormone pathway receptors and transmitters. Supplemental Table 3: Primers and probes used for functional validation.

Supplemental Data Set 1: Plant immune signalling model, version 2 (PIS-v2).

Supplemental Data Set 2: Comparison of contributions in PIS model and AtCKN subnetwork. Supplemental Data Set 3: Gene connections for network analysis results between EIN3 and NPR1.

Supplemental Data Set 4: Results of in silico regulatory element search for AtNPR1 and StNPR1.

## ACKNOWLEDGMENTS

This work was supported by grants from the Slovene Research Agency (P4-0165, J4-7636, N4-0026). We thank Katja Stare and Lidija Matičič for excellent technical assistance.

